# Biological Impact of a Large Scale Genomic Inversion that Grossly Disrupts the Relative Positions of the Origin and Terminus Loci of the Streptococcus pyogenes Chromosome

**DOI:** 10.1101/534933

**Authors:** Dragutin J. Savic, Scott V. Nguyen, Kimberly McCullor, W. Michael McShan

**Affiliations:** Department of Pharmaceutical Sciences, University of Oklahoma Health Sciences Center, Oklahoma City, Oklahoma, USA

**Keywords:** *Streptococcus pyogenes*, chromosomal rearrangements, asymmetric DNA inversions, survival, population competitions

## Abstract

A large-scale genomic inversion encompassing 0.79 Mb of the 1.816 Mb-long *Streptococcus pyogenes* serotype M49 strain NZ131 chromosome spontaneously occurs in a minor subpopulation of cells, and in this report genetic selection was used to obtain a stable lineage with this chromosomal rearrangement. This inversion, which drastically displaces the *ori* site, changes the relative length of the replication arms so that one replichore is approximately 0.41 Mb while the other is about 1.40 Mb in length. Genomic reversion to the original chromosome constellation is not observed in PCR-monitored analyses after 180 generations of growth in rich medium. As compared to the parental strain, the inversion surprisingly demonstrates a nearly identical growth pattern in exponential phase. Similarly, when cultured separately in rich medium during prolong stationary phase or in an experimental acute infection animal model (*Galleria mellonella*), the survival rate of both the parental strain and the invertant is equivalent. However, when co-incubated together, both in vitro and in vivo, the survival of the invertant declines relative to the parental strain. The accompanying aspect of the study suggests that inversions taking place nearby *ori*C, always happen to secure the linkage of *ori*C to DNA sequences responsible for chromosome partition. The biological relevance of large scale inversions is also discussed.

**IMPORTANCE:** Based on our previous work, we created to our knowledge the largest asymmetric inversion covering 43.5% of the *S. pyogenes* genome. In spite of a drastic replacement of origin of replication and the unbalanced size of replichores (1.4 Mb vs 0.41 Mb), the invertant, when not challenged with its progenitor, showed impressive vitality for growth *in vitro* and pathogenesis assays. The mutant supports the existing idea that slightly deleterious mutations can provide the setting for secondary adaptive changes. Furthermore, comparative analysis of the mutant with previously published data strongly indicate that even large genomic rearrangements survive provided that the integrity of the *ori*C and the chromosome partition cluster is preserved.

## INTRODUCTION

Circular bacterial chromosomes are replicated in a bidirectional way from the unique origin (*oriC*) and terminated at the termination region (*dif*) half a chromosome away (1, 2). The symmetry of the origin and terminus loci were shown to play the role in precise choreography of replicated chromosome separation during cell division (3). It has been shown that genetic variability depends not solely on point mutations but also to a high degree from horizontal transfer of genes and intra-genomic rearrangements, the events that may disrupt chromosome organization. Such a situation creates an “evolutionary conflict” between the tendency for stability and evolutionary drive for diversity (4). The resulting genotype represents the compromise between these two tendencies and reflects the existent selective pressure.

With the advent of genomic sequencing, the wealth of whole bacterial genomic data demonstrates that, as selective features, the distribution of genes in most cases respect the co-polarity of DNA replication and genes transcription. It also showed a strong bias towards gene activity and essentiality as well as the GC content in the leading and lagging strands (3, 5-8). Highly expressed genes and stress related genes of *Escherichia coli* and *Bacillus subtilis* were shown to be clustered on leading strands of the two opposite sides relative to the origin of replication (9). For this reason, most natural stable inversions are symmetrical at the origin-terminus axis of chromosome replication, thus maintaining gene orientation and gene distance relative to the origin of replication minimizing the disruption of the chromosome organization relative to replication and segregation (10-13).

Studies of asymmetrical inversions revealed chromosomal segments refractory to inversion. They were interpreted as deleterious effect of recombination (14) or, in contrast, as recombinational impairment of mechanical nature (15). Inversions, especially large ones, antagonize the organization of bacterial chromosome with the greatest consequences to cell viability. However, it has been shown that such rearrangements may also produce some positive trade-off for the mutants in certain extreme environmental conditions (16). We previously reported a spontaneous, large-scale rearrangement in the genome of *Streptococcus pyogenes* strain NZ131 that affects expression of the streptolysin O gene (*slo*) (17). In that study, cells with the 0.79 Mb rearranged segment of the NZ131 chromosome were a minor component of the population and could be detected solely by PCR method. No easily selectable phenotype was available to allow their isolation and study independent of the majority population. In this report, we employ genetic selection to isolate a mutant with this specific rearrangement in *S. pyogenes*, and use this large artificial inversion to assess its impact on cell viability *in vitro* and on virulence in an invertebrate acute infection model. Finally, we also performed comparative analysis of our inversion with a number of spontaneous and engineered large inversions.

## RESULTS

### Construction of the invertant OK175

An inversion-selectable mutant of *S. pyogenes* (strain NZ131, group M49) was constructed by inserting a plasmid construct harboring the distal part of the gene Spy49_*0145* and the fragment of Spy49_*1180* gene mapping 720 Kb away from the Spy49_*0145* gene and the promoter-less *erm* gene (Fig. 1a). Gene Spy49_*0145* encodes an inhibitor of the secreted streptococcal NAD-glycohydrolase, and occupies the central location on a polycistronic mRNA between the upstream NAD-glycohydrolase gene (*nga*) and the downstream streptolysin O gene (*slo*) (18). Plasmid pOK67 harboring this insert was previously described (17). At that time, the chromosomal sequence of NZ131 had not been determined so gene Spy49_*1180* was temporarily labeled as the Ex sequence, and gene Spy49_*0145* was combined into the whole *slo* upstream sequence (17). The promoter-less *ermB* gene is silent because it is preceded by the chimerical Spy49_*0145*-Spy49_*1180* sequence in which a weak *slo* promoter found inside the Spy49_*0145* gene (19, 20) is inactive due to the proximity of the Spy49_*1180* sequence (17, 20). The insertion, selected by resistance to tetracycline, occurred via recombination between the plasmid and chromosomal Spy49_*1180* sequences resulting in the strain OK173 (Fig. 1ab).

**Figure 1.**
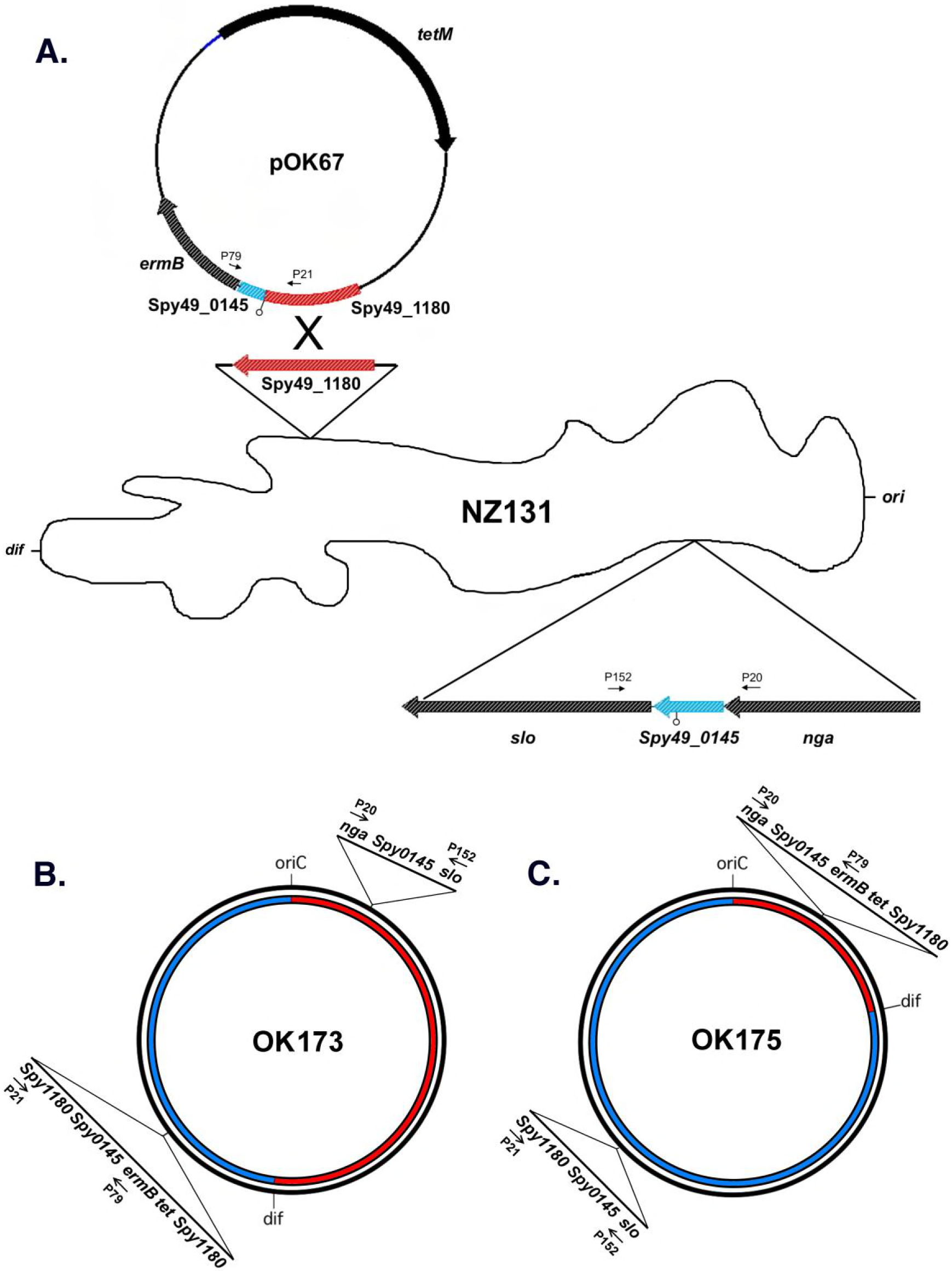
(a) Construction of the starting mutant with predilection to produce a large inversion was made by selecting integration of the plasmid pOK67 (*tet*^R^) into Spy49_*1180* gene of the strain NZ131. (b) The construct obtained (OK173) contains two inverted Spy49_*0145* gene sequences prone to recombine and generate an inversion. The origin (*ori*) and terminus (*dif*) loci are indicated (c). The generated inversion (OK175) covers 0.72 Mb (40.5%) of the approximately 1.82 Kb long *S. pyogenes* chromosome. It asymmetrically encompasses the *ori* region drastically changing its position on the chromosome. The short arrows indicate the position of PCR primers used for the verification of the constructs. The colored arcs illustrate the relative size of the replichores.

The rare inversion mutants were captured by selection on plates with erythromycin. Resultant erythromycin resistant colonies occurred following recombination between the plasmid-borne Spy49_*0145* sequences and the chromosomal Spy49_*0145* gene adjacent to *slo*, bringing transcriptionally inactive *erm* gene downstream of the strong *nga* promoter (21).

### Verification of the inversion termini

The inversion in the new construct OK175 (Fig. 1c) was verified by PCR amplification of the predicted junction sections with corresponding primers Table 2. Fig. 1. Fig. 2) and DNA sequencing of the obtained fragments (Fig. 2b). The inversion was also confirmed by PFGE of the *SgrA1* digested chromosomal DNA from both strains (Fig. 3a) and with *SmaI* and *Sfi1* (Supp. Fig. 1). Hybridization of PFGE DNA with specific origin and terminus probes pOK166 and pOK167 (Table 1) demonstrated that a compensatory rearrangement of the origin or the terminus region has not occurred (Fig. 3b).

**Table 1.** Bacterial strains and plasmids used in this study.

| Strain | Relevant characteristics | Source or reference |
| --- | --- | --- |
| *Streptococcus pyogenes* |  |  |
| NZ131 | M49 Clinical isolate genome strain | (22) |
| OK173 | Obtained by homologous recombination between chromosomal (NZ131) gene Spy49_1180* and plasmid OK67-borne fragment of the same gene (Spy49_1180). | (17) |
| OK175 | Derivative of OK173 obtained by selection for Erm<sup>R</sup> clones. Contains large inversion obtained by recombination between chromosomal and integrated pOK67-borne sequences of the Spy49_0145 gene. | This work |
| *Escherichia coli* |  |  |
| JM109 | F<sup>'</sup>/traD36 lacIq Δ(lacZ)M15 proA<sup>+</sup> B<sup>+</sup> /e14 (McrA<sup>-</sup>) Δ(lac-proAB) thi gyrA96 (Nal<sup>r</sup>) endA1 hsdR17 (r<sup>-</sup> m<sup>+</sup>) relA1 supE44 recA1 | (59) |
| **Plasmid** |  |  |
| pOK67 | Recombinant plasmid carrying 539 bp of Spy49_1180 (Ex) DNA and distal part of the gene Spy49_0145 inserted in the front of the erm gene (Erm<sup>s</sup>). | (17) |
| pOK166 | Recombinant plasmid harboring an approximately 800bp fragment from the origin region of NZ131. Obtained by cloning the product of P212 x P215 PCR reaction into the vector pGEM<sup>®</sup> T-Easy. | This work |
| pOK167 | Recombinant plasmid harboring an approximately 650bp fragment from the terminus region of NZ131. Obtained by cloning the product of P217 x P219 PCR reaction into the vector pGEM<sup>®</sup> T-Easy. | This work |
| pGEM<sup>®</sup> T-Easy | PCR cloning vector | Promega Corp. |

\* Gene as designated in the annotated NZ131 genome

**Table 2.**
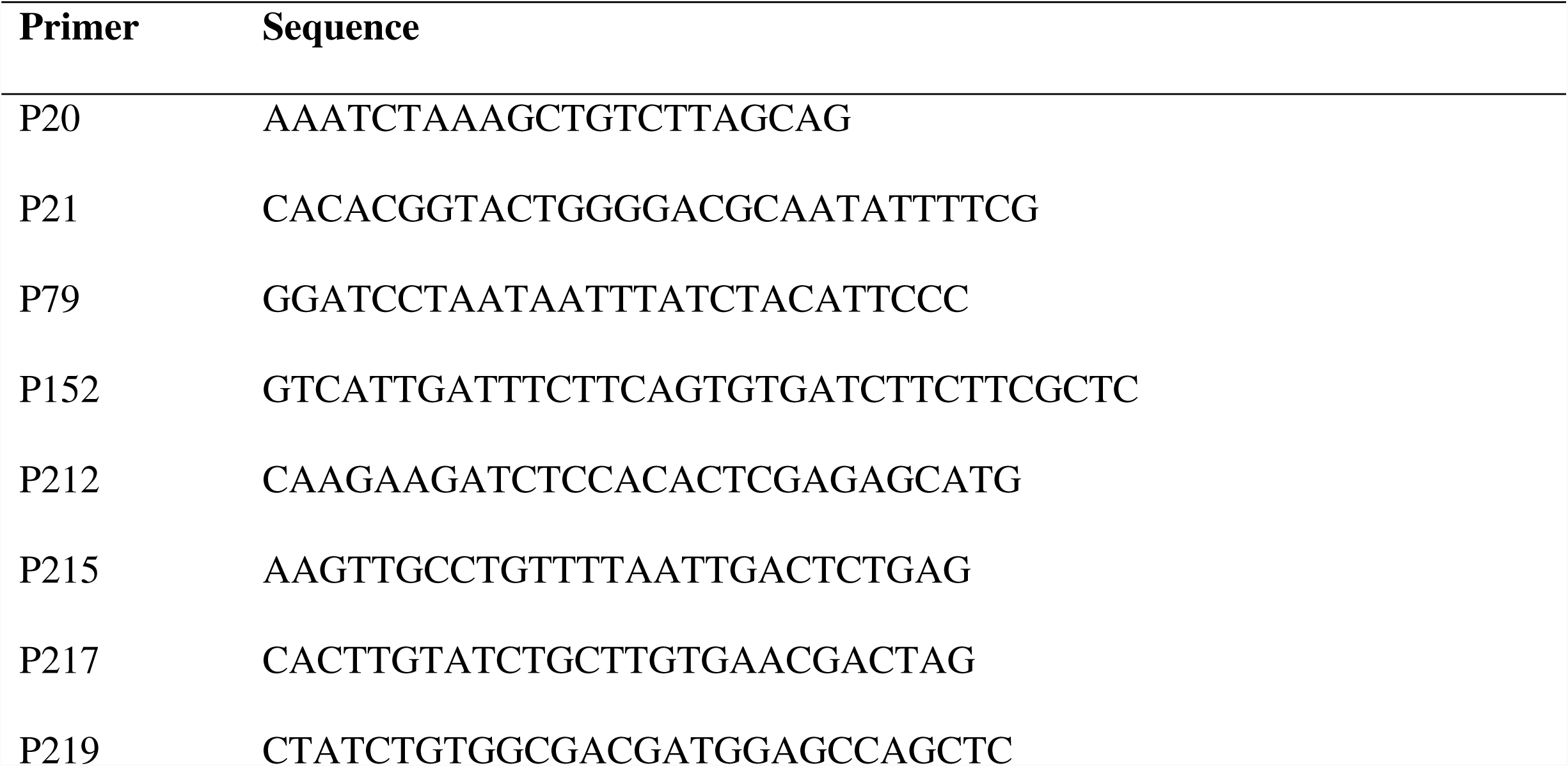
Oligonucleotide primers used in this study.

**Figure 2.**
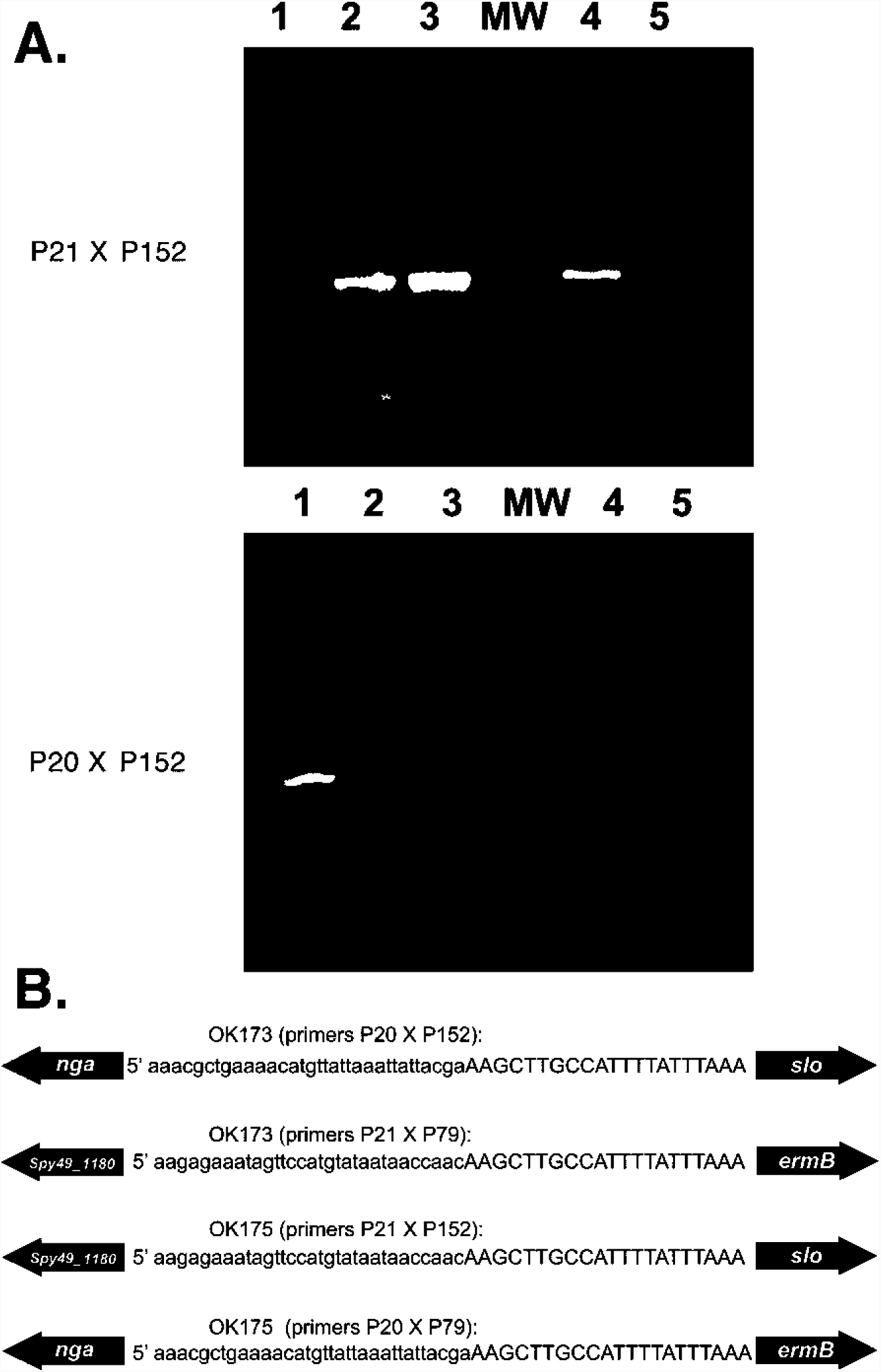
(a) PCR analysis was used to validate recombination event at the predicted regions. The primers P20, P21, P79 and P152 used in PCR reactions belong to the *nga*, Spy49_*1180, ermB* and *slo* genes respectively. PCR reactions P20 X P79 and P21 X P79 are not presented but the fragments of the expected size were obtained, purified and sequenced with other fragments. The upper part of the figure presents results of PCR reaction obtained with primers P21 X P152 while lower part shows results obtained in the reaction with P20 X P152 pair of primers. Lanes: 1) OK173; 2) OK175; 3) OK175 subcultured for 10 days; 4) Molecular size standard; 5) OK175 subcultured for 20 days. (b) Nucleotide sequences of the amplified fragments at the recombination sites in the control OK173 and the invertant OK175 strains. The nucleotides in small letters represent the *nga* gene sequence obtained from PCR fragments with primers P20 x P152 and P20 x P79, while the other two sequences (P21 x P79 and P21 x 152) are the sequence of the Spy49_*1180* gene. All of the capital letters sequences belong to the Spy49_*0145* gene.

**Figure 3.**
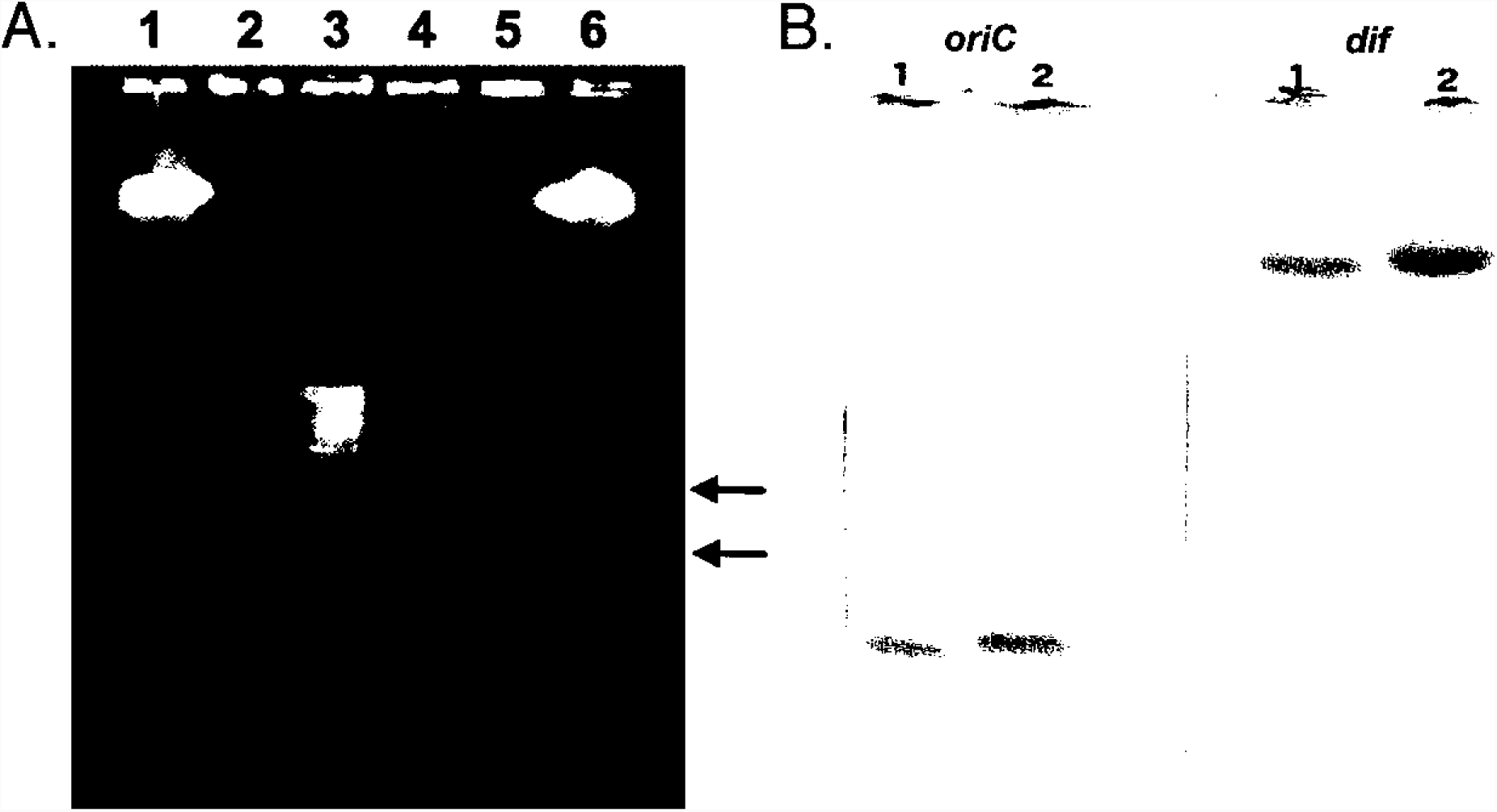
(a) PFGE of the *SgrA1* - digested chromosomal DNAs of the strains OK173 and OK175. Lanes 2 and 4 denote DNA from the preliminary strain (OK173) while lanes 3 and 5 denote DNA from the invertant (OK175). Lateral lanes are occupied with molecular weight standards. The results obtained with two other rare cutters *Sma1* and *Sfi1* were also performed (Supp. Fig. 1). Lambda DNA concatemers were used for MW standards (New England Biolabs, Ipswich, MA). (b) Southern blot of Sgr*A1* - digested DNA hybridized with probes pOK166 and pOK167 specific for *ori* and *ter* sequences. Lane 1 shows the hybridization signal obtained from digested OK173 DNA and lane 2 indicate the signal obtained from OK175 DNA.

### Testing stability of the inversion

The stability of the inversion was tested by sub culturing the strain OK175 for 20 days in THY medium. The samples taken after 10 and 20 days were diluted and 200 colonies were tested for growth on THY plates supplemented with erythromycin. Growth of all colonies suggested that the original invertant OK175 retained its original constellation after approximately 120-130 generations. The lack of any amplified DNA in the PCR reaction using P20 and P152 primers with OK175 DNA as a substrate additionally verified stability of the inversion (Fig. 2b).

### Growth patterns of NZ131, OK173 and OK175 strains

As demonstrated, the growth pattern of the invertant strain OK175 in the exponential phase does not differ much when compared to the parental strain OK173 (Fig. 4a). However, both strains display slower growth as compared to the original strain NZ131 due to the probable effect of mutation in the Spy49_*1180* gene coding for glucokinase (22). When grown in separate cultures for a prolonged period, OK173 and OK175 demonstrated similar survival rates (Fig. 4b). However, when co-cultured in a prolonged competition assay, the invertant lagged significantly behind the parental strain, and after eight days, its percentage in the mixture dropped from the initial ∼50% of the population to <0.01% in the sample of 1000 replica plated colonies (Fig. 4c).

**Figure 4.**
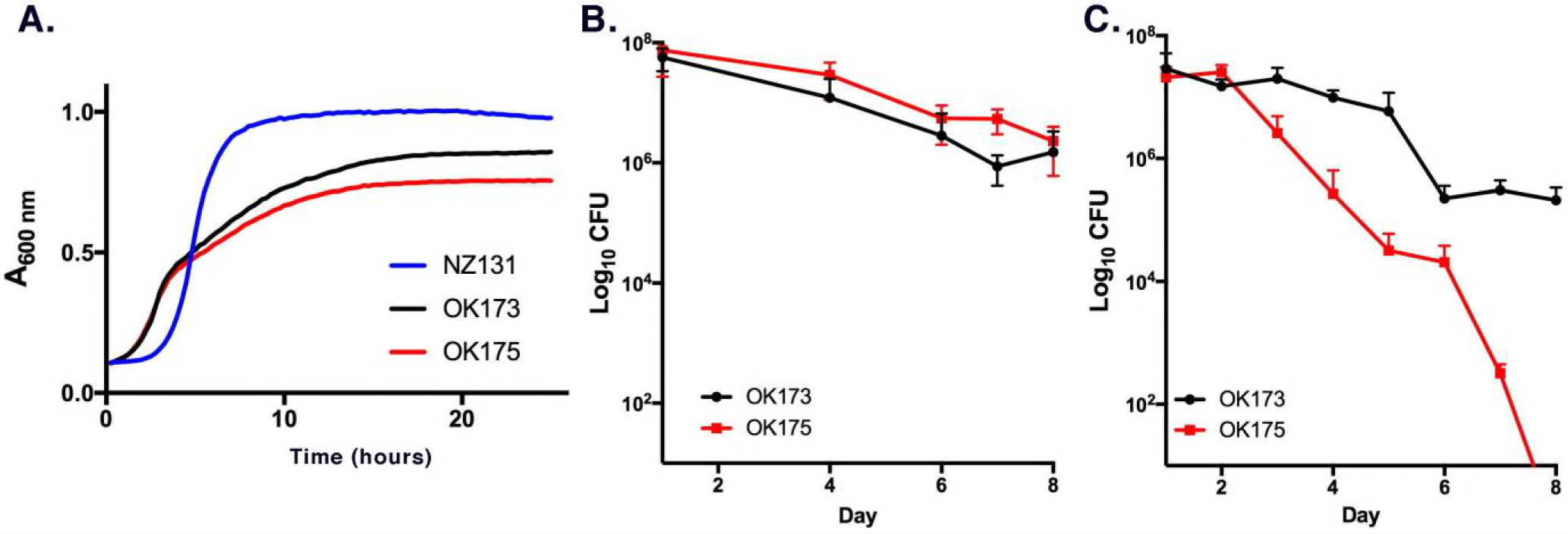
(a) The growth curves of the strains NZ131, OK173 and OK175 in the logarithmic phase. Three independents cultures were done for each strain; error bars, which were uniformly very small, are not shown for clarity of presentation. (b) The strains OK173 and OK175 grown in separate cultures in the stationary phase. (c) Reconstruction experiment of the OK173 and OK175 mixed cultures during their stationary phase. The overnight cultures of OK173 and OK175 strains were mixed of at approximately 50% ratio based on A_600_ values (Time 0). The strains were discriminated on the basis of the OK175 resistance to erythromycin.

In vivo infection of *G. mellonella* larvae with strains OK173 and OK175 demonstrated little difference in pathogenicity in monocultures (Fig. 5a). Further, when compared to the NZ131 parental strain, decreased larval mortality was observed in OK173 and OK175, possibly due to the mutation in Spy49_*1180.* Using the wax worm health index developed by Loh and co-workers (23) morbidity of the infections was also assessed (Fig. 5b), again showing no significant differences in morbidity between strain OK173 and OK175 in *G. mellonella* infections. Finally, when OK173 and OK175 were co-cultured, faster decline of OK175, as compared to the in vitro experiment (Fig. 4c) was observed (Fig. 5c).

**Figure 5.**
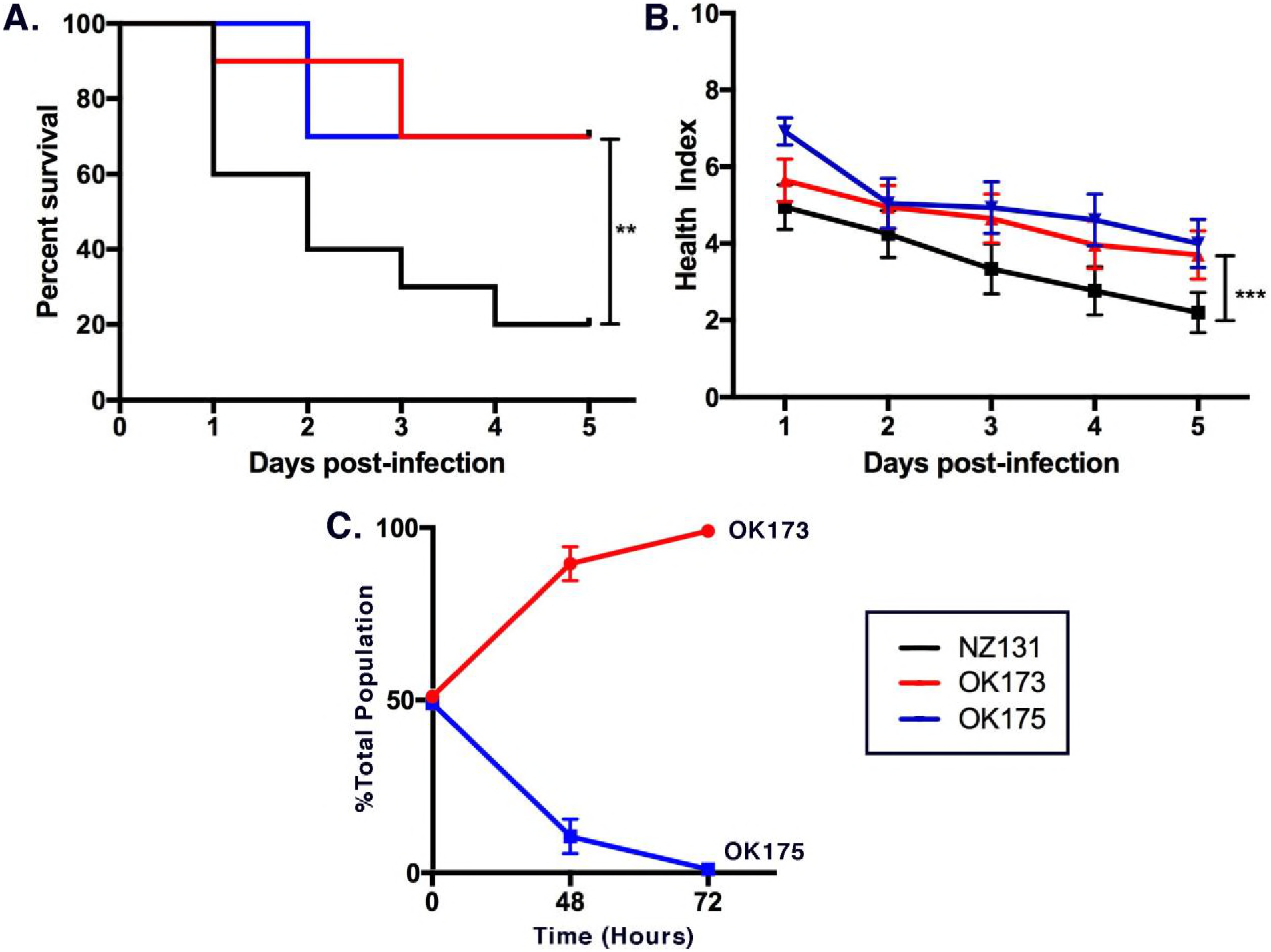
Acute infection studies. Survival (a) and mean health (b) of *G. mellonella* larvae was monitored over 96 hours post infection with NZ131, OK173, or OK175; N=10 for each group. By either metric, both OK73 and OK175 showed decreased virulence compared to parental strain NZ131. 70% of larvae challenged with either OK173 or OK175 survived after 5 days while only 20% of those infected with NZ131 did (**, *P* < 0.02; one way Mantel-Cox test). On day five, the mean health scores of larvae infected with either OK173 or OK175 were significantly different (***, *P*<0.001) from those infected with NZ131 by one-way analysis of variance (ANOVA) and Sidak’s multiple comparisons test. (c) Competition experiment with OK173 and OK175 strain.

## DISCUSSION

In the previous study, we described and characterized an unorthodox RecA-independent genomic rearrangement involving two chromosomal loci separated by 0.79 Mb on the 1.816 Mb long *S. pyogenes* chromosome (17). This rearrangement positioned a TATAAT and TCGAAA - 10 and -35 promoter sequences located in gene Spy49_*1180* 208 bp upstream from the *slo* gene. The observed rearrangement could only be detected by PCR, suggesting that only a minor subpopulation of cells carried this orientation. To demonstrate that the translocated promoter might activate *slo* gene independently of the *nga* promoter (21), we engineered the inversion by reproducing the junction between the Spy49_*1180*, the *slo* upstream sequence, and *erm*B gene as a selectable marker (Fig. 1a). However, activity of this postulated promoter was not detected (17). Finally, the inversion construct of this study which recreated the previous observation (17) failed to activate *slo* gene from the classical promoter in Spy49-*1180* gene (results not presented) (Fig. 1c). Since all experiments were performed *in vitro* (THY medium), the possibility remains that some unknown physiological factor might activate the transposed promoter or the previously identified weak *slo* promoter (19, 20) when synthesis of *slo* gene but not the *nga* gene is required.

At this point, we thought that given the size and growth characteristics, the constructed inversion deserved further study on its own right. In spite of the defect, the invertant OK175 grows at a similar rate in logarithmic phase (Fig. 4a) and survives, both *in vitro* and *in vivo*, similarly in a protracted stationary phase when compared to the parental mutant OK173 (Fig. 4b, Fig. 5ab). However, the situation changes when OK173 and OK175 are cultured together (Fig. 4c Fig. 5c). OK175 has diminished fitness when co-cultured with OK173 in depleted medium and scavenging for the remaining nutrients. Additionally, with the *in vivo* waxworm model, OK175 is outcompeted by OK173 in co-cultures much faster that may be due to the Slo^-^ phenotype of OK175 (Fig. 5c) These results show great similarity to the previous study which demonstrated that in naturally non-transformable *E. coli,* extracellular DNA can serve as the sole source of carbon and energy in nutrient-starved conditions (24). Genome sequencing revealed that horizontal gene transfer is frequent among strains of *S. pyogenes,* and it is reasonable to propose that *in vivo* DNA transfer may occur by mechanisms other than transduction as has been seen in *E. coli* (22, 25). In their seminal work, Mashburn-Warren and collaborators demonstrated that in *S. pyogenes* all functional genes necessary for transformation exist, and while the pathway does not typically function in laboratory media it can occur in biofilms (26, 27). Thus, it is realistic to speculate that *S. pyogenes*, similarly to *E. coli* (24), is also in possession of an additional double-stranded DNA uptake system that is functional to some degree in OK173 but not in the invertant OK175 that consequently is unable to compete with OK173 for scavenging DNA in depleted medium (Fig. 4c).

In contrast to spontaneous, X-shaped inversions in which the rearrangements reverse the genomic sequence symmetrically around the origin of replication (10, 13, 28) the inversion we report here drastically displaces the *ori* site, resulting in a dramatic change of replichore sizes from both having approximately the same lengths to 0.41 Mb and 1.40 Mb, respectively. The terminus region has not been closely studied in strain NZ131, but its location opposite the origin of replication, as expected, can be easily deduced from the switch of the GC content and polarity of transcription in leading and lagging strands as well nucleotide comparative analysis (22)(see Supplemental Fig. 2).

Bacterial chromosomes are adaptively organized to optimize cellular functions such as chromosome replication, segregation and resolution, symmetrical polarity of transcription, gene dosage (including optimal positioning of highly expressed and stress genes), and coordinated gene expression (4). All of these functions should have been affected to some degree in the OK175 invertant. They obviously are but given the size and eccentricity of the inversion on one side, and stability and almost normal growth rate of the invertant, these effects are surprisingly not considerably pronounced. A facultative pathogen, *S. pyogenes* shows many repeated elements like phage chromosomes, ribosomal operons and IS elements (13, 16, 22) resulting in frequent chromosome rearrangements that allow fast adaptation by permitting generation of antigen variations. As speculated before, *S. pyogenes* and similar organisms containing such recombination elements, as is disclosed in frequent rearrangements, have possibly relaxed the selection of some chromosomal features of their chromosomes (4). Also, both strains OK173 and OK175 demonstrate, possibly due to the mutated Spy49_*1180* gene, slower growth relative to NZ131 (Fig. 1a), which may attenuate the harmful effect of the inversion (29).

It is intriguing to compare inversion of this study with large inversions described in the extensive study of the engineered inversions in closely related low G+C member of Streptococcaceae family Firmicute, *Lactococcus lactis* (29). Of the strains described in that report, the one having a large asymmetric inversion encompassing *oriC* (strain *pcp-12;* 44% of the chromosome inverted) is similar to OK175 insomuch as the inversion is stable for up to 150 generations. By contrast, the two other *L. lactis* strains with large asymmetric inversions in the same region (*pcp-6* and *pcp-31*), which are shorter than *pcp-12* and have end points closer to *oriC*, are less stable. The longer of these two (*pcp-31)* is stable in synthetic medium but unstable in rich medium, and the one with the smaller inversion (*pcp-6*), is unstable in both media (29).

Studies on *Bacillus subtilis*, demonstrate that bacterial cells, similarly to eukaryotic cells, use a mitotic-like mechanism (Par) to provide accurate chromosome partitioning and separation of the replicated sister replication origins. These results identified the *parABS* genes, which are almost invariably located close to the origin of chromosome replication (*oriC*). Chromosomal centromere-like *parS* sequences flank the origin of replication (30, 31). ParB is a site-specific DNA-binding protein (SpoOJ in *B. subtilis*) that binds *parS* sites (31-33) and then spreads laterally along the DNA, perhaps for several kb (34-36). It also controls the product of *parA,* which affects initiation of DNA replication (37). However, the main role of ParB (SpoOJ) is its role in recruitment of the SMC proteins to the *oriC* site, the proteins being conserved from bacteria to humans. Most bacteria encode a single SMC protein that forms a homodimer (38). The fact that SMC complexes are highly enriched at the region around the origin of replication indicates that they are directly involved in proper segregation (39-42). The conserved positional relationship of *parS, parB, parS and oriC* suggests that these genes have coevolved and that there is a selective pressure promoting their proximity. All these data provide support for the notion that bacterial cells use a mitotic-like mechanism to provide accurate chromosome partitioning, separation of the replicated sister origins and regulation in *B. subtillis* with crucial role of SpoOJ (ParB) and to a lesser degree Soj (ParA) (43) and their orthologs in other bacteria (44).

Considering the clustering of partition relevant genes around the *oriC,* one can propose that the described largest *pcp-12* inversion, as compared to the smaller *pcp-6* and *pct-31* inversions, of *L. lactis* (29), is the one that does not disrupt partition complex around *oriC* but translocates it *en block* to the different location of the chromosome. In an attempt to identify *parS* sites in *S. pyogenes* chromosome, we initially used a BLASTN search with the *parS* consensus sequence TGTTNCACGTGAAACA from in *B. subtilis* (31) but with no success. However, the *parS* consensus sequence from *S. pneumoniae* genome (tGTTTCACGtGAAACa) (45) identified three hits in immediate vicinity of the *oriC*: TGTTTCACGTGAA (from 1209– 1221 bp); TGTTTCACGTGA (1293-1304 bp) and TCACGTGAAAC (2272-2282 bp). The first two sequences in *S. pyogenes* are direct repeats that form inverted repeats to the third sequence. In addition, a *parB* gene was identified at the very end of the NZ131 chromosome in the immediate proximity to *oriC* (22). Thus, the predicted spindle formation apparatus in *S. pyogenes* appears to be highly clustered around the origin. Based upon this inference, the inversion in OK175 strain, which is similar to the inversion strain *pcp-12* of *L. lactis,* translocates the whole *parBC* cluster to a new distant location on the chromosome. In addition, the most recent study demonstrated that the genome of the hyper virulent strain of *S. pyogenes* M23ND underwent complex rearrangements that included multiple DNA translocations, inversions and integrations (16). In one inversion, found in six independent isolates, the *ori*C– proximal recombination site was found to be located only 3 Kb counterclockwise of the *par*B site ensuring the linkage of that gene to the *ori*C keeping in that way the partition complex intact. Finally, in the long-term evolutionary experiment (25 years) with *E. coli,* an inversion covering approximately 1.5 Kb of the chromosome, was isolated (46). Again, the *ori*C proximal recombination site occurred approximately 0.28 Mb counterclockwise of *ori*C keeping the nearby sequences linked to the *ori*C. Finally, it was reported that MukBEF, a member of the SMC superfamily is preferentially localized within the region of at least 0.4 Kb centered on *ori*C (41) and that recruitment of MukBEF towards the origin region is essential for chromosome segregation in *E. coli* (47). Thus, the unforeseen aspect of this study, comprehended by comparative analysis of the OK175 invertant and the invertant mutants reported above, suggest that optimal functioning of the spindle apparatus requires strict clustering of their genetic determinants in the proximity of *ori*C and that their *en bloc* transposition, by inversions, and possibly by other types of rearrangements, even to the distant part of the chromosome, does not necessarily have immediate consequences to the cell. As cited before, colocalization of *ori*C and genes responsible for chromosome partition had been well established, but their obligatory linkage in the course of chromosomal rearrangements, to our knowledge, had never been discussed.

The results of this work show that *S. pyogenes* can also tolerate, at least for some time, drastic chromosomal rearrangements. This inversion, and probably similar chromosomal perturbations, are certainly not adaptive for long-term survival *in vivo*, but one could imagine that under strong environmental stress, they might serve as a raw genetic material for further adaptive chromosome rearrangements. As demonstrated by theoretical models with higher organisms, small populations under stress followed by random drift can promote the accumulation of not only neutral mutations but also of slightly deleterious mutations, in which the resulting alterations to chromosome structure can provide potential settings for secondary adaptive changes that are not possible in large populations (48, 49). In our experimental system, extended growth in mixed and single cultures simulates both stress and genetic drift. It also mimics small populations that in higher organisms lack or have a very low incidence of competition. All of these factors are present in the mixed cultures resulting in the elimination of the mutant. During infection, *S. pyogenes* certainly experiences stressful conditions accompanied with reduction in population size, the situation in which the magnitude of selection intensity can be overwhelmed by the stochastic force of random genetic drift (48). Further inquiries along this line may be of interest in studies of streptococcal pathogenesis.

## MATERIAL AND METHODS

### Bacterial strains, media and culture

Bacterial strains and plasmids used in this study are described in Table 1. *S. pyogenes* strains were grown in Difco Todd-Hewitt medium containing 0.2% yeast extract (THY). When needed, 1.5 µg/ml of erythromycin and 12.5 µg/ml of tetracycline were added to the plates. *Escherichia coli* was grown in Luria-Bertani (LB) medium. When required, 500 µg/ml of erythromycin and 12.5 µg/ml of tetracycline were added to the plates. To make plates, both THY and LB media were solidified with 1.5% agar. The growth of bacterial cultures in the exponential phase was monitored using the Bioscreen-C automated microbiology growth curve analysis system (Growth Curves USA, Piscataway, NJ). For quantifying CFU, the described method from Jett *et al.* was employed (50). When grown in the mixed long-term stationary cultures, the strains were discriminated on the basis of their sensitivity or resistance to erythromycin.

### DNA manipulation, isolation, and sequencing

Plasmid isolation, digestions with restriction enzymes, ligation, Southern Blot and other recombinant DNA techniques were performed by standard procedures (51). Plasmids were introduced into *S. pyogenes* recipients by electrotransformation (52), and transformation of *E. coli* was performed by standard CaCl_2_ method (53). Plasmids were maintained in *E. coli* host JM109 and isolated with a commercial plasmid isolation kit (Qiagen Inc.). For cloning PCR fragments, specialized cloning vectors with T-nucleotide overhangs pT7Blue (Novagen) and pGEM-T Easy (Promega) were used. All sequencing reactions were performed at the University of Oklahoma Health Sciences Center Laboratory for Genomics and Bioinformatics core sequencing facility.

### PCR analysis

PCR oligonucleotide primers used in this study are listed in Table 2 and were purchased from Integrated DNA Technologies (Coralville, IA). Streptococcal template DNA was isolated by the guanidium thiocyanate DNA isolation method (54). In most cases 40 cycles of amplification were carried out in a Techne DNA thermocycler with the following cycle: strand denaturation (1min at 94° C), annealing (1 min at 59° C) and elongation (1 min at 72° C). The total volume of PCR mixture was 50 µl, and consisted of Taq DNA polymerase (1 U), Taq polymerase buffer (1 X), Mg^2+^ (2mM), deoxynucleotide phosphates (200 µM each) and primers (1 µM each). Following amplification, 10 µl of each reaction was analyzed by agarose gel electrophoresis. When used for DNA sequencing, PCR products were purified with PCR purification kit following the manufacturer’s protocol (Qiagen, Inc.).

### Pulse field electrophoresis

CHEF gel was performed as previously described (55) with some modifications. A portion of the agarose plug containing total DNA from each strain was excised and digested with either *SgrA1, SmaI,* or *Sfi1* restriction endonucleases at 37°C for 16 h. The plugs were washed in 1 ml 10 mM Tris-1mM EDTA, pH 8 for 1 h at 37°C and then loaded into the wells of 1% pulse-field certified agarose gel (Bio-Rad Laboratories) in 0.5X TBE (45mM Tris HCl, 45mM boric acid, 1 mM EDTA). The digested DNA was separated by electrophoresis using a CHEF DRII device (Bio-Rad Laboratories) with a pulse ramped from 5 to 35 sec for 21 hr. at 200 V. The gels were then stained with ethidium bromide and photographed before DNA fragments were transferred to membranes by capillary blotting. As described below, the membranes were further used for hybridization with molecular probes harboring chromosomal origin and terminus sequences.

### Construction of molecular probes for detecting chromosomal restriction fragments with origin and terminus sequences

The probes that contained the sequences around origin and terminus of replication were amplified from these regions of the NZ131 genome sequence (22). The origin of replication for *S. pyogenes* has been previously reported (56), and the terminus was deduced by homology with known *dif* sequences (Supp. Fig. 2). DNA probes specific for these sites were prepared using primers P212 and P215 and P217 and P219 (Fig. 2) for the origin and terminus regions respectively. The PCR products were labeled with digoxigenin-11-dUTP (Roche Diagnostics) and done according to the manufacturer’s recommended conditions. Hybridization, stringency washes, and detection of the bound probe were also performed according to the manufacturer’s recommended protocol and reagents.

### *Galleria mellonella* acute infection in vivo model

Larvae of the greater wax moth *Galleria mellonella* (commonly referred to as wax worms) were reared in-house in a dedicated 37°C incubator and fed an antibiotic-free, artificial diet of 227g of Gerber mixed-grain baby cereal, 80 ml honey, and 80 ml glycerin as previously described (57). Larvae were selected at the last instar before pupation and then kept at 4°C for 15 minutes before injection. Overnight *S. pyogenes* cultures were grown in THY broth at 37°C and harvested by centrifugation at 5,000 x g for 5 minutes then resuspended in standard phosphate buffered solution (PBS) to achieve a desired concentration between 1 x 10^7^ - 2 x 10^7^ CFU/ml. Suspensions were standardized at OD_600_ nm (OD_600_ nm between 0.5-0.6) in PBS to achieve the final concentration. Intrahaemocoelic injection of the larvae (n=10 per condition) was performed by a 1 mL tuberculin syringe (Norm-Ject) fitted with a custom 30½-gauge needle (BD PrecisionGlide) at a volume of 20 µL (∼20,000 cells) and using a KDS 100 automated syringe pump (KD Scientific, Holliston, MA). Blank PBS (20 µL) was injected into an equal number of control larvae. Larvae were injected at the hindmost left proleg as previously described (23, 58). Post injection larvae were incubated in 9 cm petri dishes at 37°C for 5 days. Worm health index and mortality were scored as described previously (23).

### *Galleria mellonella* competition assay

Overnight cultures of *S. pyogenes* strains OK173 (erythromycin sensitive) and OK175 (erythromycin resistant) were prepared as described above. Cultures were then mixed and resuspended to achieve a 50:50 ratio. Serial dilutions of this preparation were plated on sheep’s blood agar plates and on THY agar plates supplemented with 3 µg erythromycin/ml to determine culture ratios. 20 µL of this suspension was injected into worms as described above. Larvae were injected (n = 15) and incubated at 37°C in a 9 cm Petri dish. At intervals of 24, 48, 72, 96, and 120 hours post-infection (PI), groups of 3 worms were chosen at random and sacrificed. Each worm was homogenized by mechanical disruption and resuspended in 0.5 ml PBS as previously described (23). Aliquots of this suspension were serially diluted and plated as previously described to determine the relative CFUs of OK173 and OK175 post-infection.

## ACKNOWLEDGEMENTS

We thank John R. Roth and N. Tucic† for the help in interpreting some experimental data. We also thank Catherine J. King and Arto S. Baghdayan for expert technical assistance. This work was made possible by an Oklahoma Center for the Advancement of Science and Technology (OCAST) grant HR11-133 to WMM.

---

**Supplemental Figure 1.**
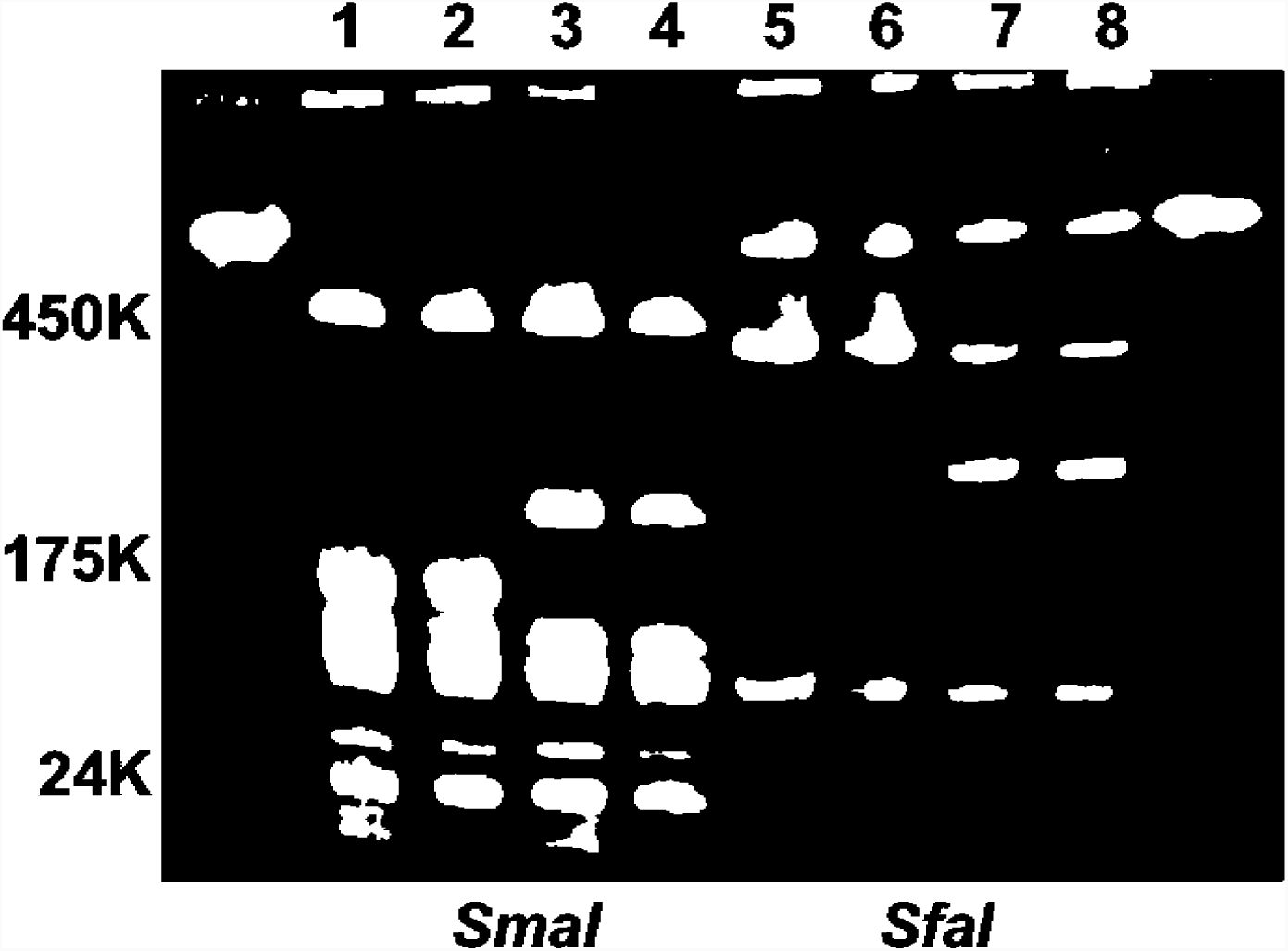
PFGE of chromosomal DNAs from strains NG131, OK173 and OK175 digested with rare cutter restriction endonucleases *Sfi1* and *SgrA1*. Lanes 1 and 5 denote DNA from NZ131, lanes 2 and 6 represent DNA from the OK173, lane 7 shows DNA from the invertant OK175 and lane 8 represent DNA from OK175 grown for 150 generations.

**Supplemental Fig. 2.**
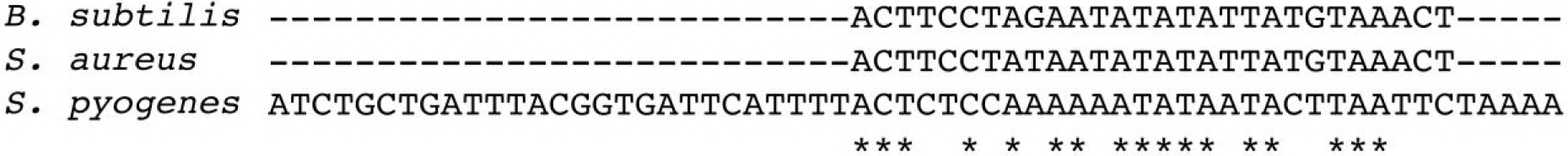
The predicted replication terminus site (*dif*) was identified by BLASTN alignment of the *S. pyogenes* genome with the known *dif* sites from *Staphylococcus aureus* and *Bacillus subtilis* (60).

